# Enhanced axonal Neuregulin-1 type-III signaling ameliorates disease severity in a CMT1B mouse model

**DOI:** 10.1101/386730

**Authors:** Cristina Scapin, Cinzia Ferri, Emanuela Pettinato, Desiree Zambroni, Francesca Bianchi, Ubaldo Del Carro, Sophie Belin, Donatella Caruso, Nico Mitro, Marta Pellegatta, Carla Taveggia, Markus H. Schwab, Klaus-Armin Nave, M. Laura Feltri, Lawrence Wrabetz, Maurizio D’Antonio

## Abstract

Charcot–Marie–Tooth neuropathies (CMTs) are a group of genetic disorders that affect the peripheral nervous system (PNS) with heterogeneous pathogenesis and no available treatment. Axonal Neuregulin 1 type III (Nrg1TIII) drives peripheral nerve myelination by activating downstream signaling pathways such as PI3K/Akt and MAPK/Erk that converge on master transcriptional regulators of myelin genes, such as Krox20. We reasoned that modulating Nrg1TIII activity may constitute a general therapeutic strategy to treat CMTs that are characterized by reduced levels of myelination. Here, we show that genetic overexpression of Nrg1TIII ameliorates neurophysiological and morphological parameters in a mouse model of demyelinating CMT1B, without exacerbating the toxic gain of function that underlies the neuropathy. Intriguingly, the mechanism appears not to be related to Krox20 or myelin gene upregulation, but rather to a beneficial rebalancing in the stoichiometry of myelin lipids and proteins. Finally, we provide proof of principle that stimulating Nrg1TIII signaling, by pharmacological suppression of the Nrg1TIII inhibitor TACE/ADAM17, also ameliorates the neuropathy. Thus, modulation of Nrg1TIII by TACE/ADAM17 inhibition may represent a general treatment for hypomyelinating neuropathies.

## INTRODUCTION

Myelin is an electrically insulating sheath formed in the peripheral nervous system (PNS) by a protrusion of the Schwann cell plasma membrane, which enwraps axons facilitating nerve impulse transmission and supporting long term integrity of axons (Jessen & Mirsky, 2005, Nave, 2010). Charcot-Marie-Tooth (CMT) diseases are the most common de- and dys-myelinating hereditary neuropathies, with onset in childhood or early adulthood. CMTs are characterized by muscular weakness, tremor, hind-limbs atrophy and foot deformities; at present there is no treatment (Barisic, Claeys et al., 2008, Reilly & Shy, 2009, Scherer & Wrabetz, 2008). CMTs are due to mutations in at least 70 different genes, and although clinical and pathological features partially overlap amongst CMT subtypes, the analysis of the molecular mechanism has revealed a highly heterogeneous pathogenesis (Rossor, Polke et al., 2013). As such, it would be very useful to identify unifying therapeutic strategies. Most CMT neuropathies display altered levels of myelination and, therefore, the development of strategies aimed at restoring normal levels of myelin could result in an effective approach for their collective treatment.

Nrg1TIII, a member of the NRG1 family of proteins, is an essential instructive signal in peripheral myelination. Axonal Nrg1TIII drives a Schwann cell binary choice between a myelinating or non-myelinating phenotype (Taveggia, Zanazzi et al., 2005), and the levels of Nrg1TIII determine the amount of myelin and its thickness (Michailov, Sereda et al., 2004, Taveggia et al., 2005). Nrg1TIII mostly acts through signaling pathways that control master regulators such as the transcription factor Krox20 (Egr2 gene), which activates genes for the synthesis of myelin proteins and lipids (Leblanc, Srinivasan et al., 2005, Parkinson, Bhaskaran et al., 2004). Nrg1TIII activity is regulated by proteolytic shedding: the β-secretase BACE1 cleaves Nrg1 and positively regulates myelination (Hu, Hicks et al., 2006, Willem, Garratt et al., 2006), whereas the α-secretase TACE cleaves Nrg1TIII to inhibit myelination (La Marca, Cerri et al., 2011). Importantly, recent work has indicated that genetic overexpression or soluble administration of Nrg1 overcomes impaired nerve development in rodent models of CMT1A (Fledrich, Stassart et al., 2014).

Amongst CMTs, those due to Myelin Protein Zero (*MPZ*) mutations (CMT1B) are relatively common (Scherer & Wrabetz, 2008, Shy, 2006). *MPZ* encodes for P0, the most abundant glycoprotein of peripheral myelin (D’Urso, Brophy et al., 1990, Shapiro, Doyle et al., 1996). At least 200 mutations in *MPZ* have been identified and the phenotype varies widely from mild to more severe forms (Sanmaneechai, Feely et al., 2015, Shy, 2006, Timmerman, Strickland et al., 2014). We have generated a series of transgenic mouse models of CMT1B that present various degrees of hypomyelination, demyelination and inefficient remyelination of peripheral nerves and shown that the molecular mechanisms include both loss and gain of toxic function (Wrabetz, D’Antonio et al., 2006). While it is reasonable to expect that augmenting Nrg1 signaling rescues neuropathies due to P0 loss of function, it is important to understand if Nrg1 signaling can also overcome the intoxicated pathways and ameliorate neuropathy when the mutations determine a complex gain of function scenario. This is very relevant since in humans pure P0 loss of function mutations are very rare, and for most mutations the mechanism includes toxic gain of function. For example, deletion of serine 63 in the extracellular domain of P0 (P0S63del) in transgenic mice causes a CMT1B like neuropathy that mimics the corresponding human disease (Wrabetz et al., 2006). In normal condition P0 is targeted from the synthetic pathway to the myelin sheath (Trapp, Itoyama et al., 1981, Trapp, Kidd et al., 1995). In contrast, P0S63del is misfolded and almost completely retained in the endoplasmic reticulum (ER), where it elicits an unfolded protein response (UPR) (D’Antonio, Musner et al., 2013, Pennuto, Tinelli et al., 2008, Wrabetz et al., 2006). Importantly, intracellular retention of mutant proteins has been described for CMT1B (Bai, Wu et al., 2018) and for several other CMT disease genes (for example PMP22 and Cx32) and some of these also cause a UPR (Deschenes, Walcott et al., 1997, Giambonini-Brugnoli, Buchstaller et al., 2005, Okamoto, Pehlivan et al., 2013, Yum, Kleopa et al., 2002).

Here we show that genetic or pharmacological modulation of Nrg1TIII activity improves hypomyelination in the P0S63del model, without exacerbating the ER stress due to the toxic gain of mutant P0 function. Surprisingly, this improvement is independent of Krox20 regulation. Our data open new prospects for the identification of a therapy for hypomyelinating CMTs.

## MATERIAL AND METHODS

### Transgenic mice

All experiments involving animals were performed in accordance with experimental protocols approved by the San Raffaele Scientific Institute Animal Care and Use Committee. P0S63del mice and their genotype analysis were previously described (Wrabetz *et al.*, 2006). HANI transgenic mice (Velanac, Unterbarnscheidt et al., 2012) were generated by insertion of a transgene containing Nrg1TIII fused to a N-terminal HA-tag under a constitutive neuronal Thy 1.2 promoter (Caroni, 1997). P0S63del mice were maintained on the FVB/N background, whereas HANI mice were on the C57B6/N genetic background. The analysis of WT, HANI/+, S63del or HANI/+//S63del mice was performed in F1 hybrid background FVB//C57B/6. In all the experiments, littermates were used as controls.

### Myelinating Dorsal Root Ganglion (DRG) explant cultures and immunohistochemistry analysis

Mouse DRG explants were isolated from E13.5 WT and S63del embryos, seeded on collagen-coated plates and maintained in culture as described (Taveggia et al., 2005). Myelination was induced with 50μg/ml ascorbic acid (Sigma-Aldrich). Pharmacological treatment with BMS-561392 (Bristol Myers Squibb), reconstituted in DMSO, was performed for 10 days in parallel to the induction of myelination. Immunofluorescent staining on myelinated DRG explant cultures was performed as previously described (D’Antonio et al., 2013). The following antibodies were used: chicken anti Neurofilament (1:1000; BioLegend) and rat anti MBP (smi99/94; 1:5; BioLegend), FITC- and Rhodamine-coniugated (1:200; Jackson ImmunoResearch, Baltimore, MD, USA). The number of MBP-positive internodes was counted from 10 fields per DRG, from 3 DRG cultures per genotype. Immunofluorescence was imaged with a Leica DM5000 microscope, and images were acquired with a Leica DFC480 digital camera and processed with Adobe Photoshop CS4 (Adobe Systems, San Jose, CA).

### Western blot analysis

Sciatic nerves from transgenic mice were dissected and frozen in liquid nitrogen. Proteins extraction and protein content analysis were previously described (Wrabetz, Feltri et al., 2000). Rabbit polyclonal antibodies recognized MAG (1:1000; Invitrogen), PMP22 (1:2000; Abcam), PMP2 (1:1000; Abcam), Krox20 (1:1000; Covance), Akt and P-Akt^(Ser473)^ (1:1000; Cell Signaling), Erk1/2 and P-Erk1/2 (1:1000; Cell Signaling), Grp78/Bip (1:1000; Novus Biological), Calnexin (1:2000; Sigma Aldrich) and Nrg1 C-terminal domain (1:500; Santa Cruz Biotechnology). Rabbit monoclonal antibodies recognized eIF2α and P-eIF2α (1:2000; Cell Signaling XP-Technology). Rat monoclonal antibodies recognized Grp94 (1:2000; Abcam) and HA-tag (1:2000; Roche). Chicken monoclonal recognized P0 (1:2000; Aves). Mouse monoclonal recognized MBP (1:5000; Covance) and ß-tubulin (1:5000; Sigma). Peroxidase-conjugated secondary antibodies (1:3000; Sigma Aldrich) were visualized using the ECL method with autoradiography film (GE Healthcare). Densitometric quantification was performed with ImageJ.

### Electrophysiological analyses

The electrophysiological evaluation was performed on 8-10 mice/genotype at 6 months of age with a specific EMG system (NeuroMep Micro, Neurosoft, Russia), as previously described (Biffi, De Palma et al., 2004). Mice were anesthetized with trichloroethanol, 0.02 ml/g of body weight (25-30 gr), and placed under a heating lamp to maintain constant body temperature. Sciatic nerve conduction velocity (NCV) was obtained by stimulating the nerve with steel monopolar needle electrodes. A pair of stimulating electrodes was inserted subcutaneously near the nerve at the ankle. A second pair of electrodes was placed at the sciatic notch to obtain two distinct sites of stimulation, proximal and distal along the nerve. Compound motor action potential (cMAP), was recorded with a pair of needle electrodes; the active electrode was inserted in muscles in the middle of the paw, whereas the reference was placed in the skin between the first and second digit. Sciatic nerve F-wave latency measurement was obtained by stimulating the nerve at the ankle and recording the responses in the paw muscle, with the same electrodes employed for the NCV study.

### Morphological and morphometric analyses

Transgenic and control littermates were sacrificed at the ages indicated and sciatic nerves were dissected. Semi-thin section and electron microscope analyses of sciatic nerves, at P28 and 6 months, were performed as previously described (Quattrini, Previtali et al., 1996). The number of demyelinated axons and onion bulbs were counted blind to genotype from 6-month old sciatic nerve semi-thin sections (0.5-1 μm thick) stained with toluidine blue; images were taken with a 100x objective. 800–1600 fibers in 10–20 fields for each animal in 3 nerves per genotype were analyzed. G-ratio analysis (axonal diameter/fiber diameter) and distribution of fiber diameters for myelinated axons were performed with a semi-automated system (Leica QWin V3) as described (D’Antonio et al., 2013, Pennuto et al., 2008). 10–15 microscopic fields from nerves of 3–5 animals per genotype at each timepoint were analyzed.

### Immuno-electron microscopy (IEM)

IEM was performed on transgenic and control nerves at P30. After left ventricular perfusion with 4% paraformaldehyde/2.5% glutaraldehyde in 0.08 M sodium phosphate buffer (pH 7.3), sciatic nerves were dissected and fixed in the same solution overnight at 4°C. Samples were infiltrated with increasing gradients (1M, 1.5M, 2M, 2.3M) of sucrose-30% polyvynilpyrrolidone (PVP) solution for 24 hours for each incubation with constant agitation, and then stored at 4°C in 2.3M sucrose/ 30% PVP for three days before cutting. Tissues were mounted on specimen pins and frozen in liquid nitrogen. Ultrathin cryosections (60nm thick) were sectioned using an ultracut ultramicrotome and processed as described previously (Trapp, Andrews et al., 1989, Yin, Kidd et al., 2000) with the following modifications: the cryosections were collected over nickel formvar coated grids and treated with gelatin 2% and PGB (0.1M glycine, 1% BSA, in PBS 1X) and stained with primary antibody (chicken P0, Aves, 1:600 in PGB) for 1 hour at 37°C. Sections were then rinsed in PGB and incubated with secondary antibody (1:200 rabbit anti chicken – 10nm gold particles conjugated-British Biocell International) for 1 hour at room temperature. After rinsing in cacodylate buffer 0.12M, samples were incubated in osmium 1% in cacodylate buffer 0.12M and then stained in saturated uranyl acetate. Tissues were then rinsed in water, dehydrated in increasing gradients of ethanol and finally incubated in ethanol 100%-LR White 1:1 and then in LR white quickly, and the grids were left overnight at 60°C. Cryosections were then stained with saturated uranyl acetate and lead citrate and examined by EM.

### TaqMan^®^ quantitative polymerase chain reaction analysis

Sciatic nerves from WT and transgenic mice were frozen in liquid nitrogen after dissection. Total RNA from at least 3 sciatic nerves (P28 or 6 months) for each genotype was extracted using TRIzol (Roche Diagnostic GmbH, Germany) and reverse transcription was performed as described previously (D’Antonio et al., 2013, Wrabetz et al., 2006). Quantitative PCR was performed according to manufacturer’s instructions (TaqMan, PE Applied Biosystems Instruments) on an ABI PRISM 7700 sequence detection system (Applied Biosystems Instruments). The relative standard curve method was applied using WT mice as reference. Normalization was performed using 18S rRNA as reference gene. Target and reference gene PCR amplification were performed in separate tubes with Assay on Demand^™^ (Applied Biosystems Instruments): 18S assay Hs99999901_s1; MPZ assay Mm00485139_m1; MAG assay Mm00487541_m1; PMP22 assay Mm01333393_m1; MBP assay Rn00690431_m1; Ddit3/Chop, Mm00492097_m1; Xbp-1u assay, Mm00457357_m1; Xbp-1s assay, Mm03464496_m1; Hspa5/BiP assay, Mm00517691_m1; Egr2/Krox20 assay, Mm00456650_m1. To quantify the relative abundance of WT and mutant P0 we designed the following allelic discrimination assays: P0wt assay forward TGCTCCTTCTGGTCCAGTGAAT, reverse GGAAGATCGAAATGGCATCTCT and probe Mgb ATGACATCTCTTTTACCTGG; P0mut assay forward TGCTCCTTCTGGTCCAGTGAAT, reverse GGAAGATCGAAATGGCATCTCT and probe Mgb ATGACATCTTTACCTGGC.

### Lipidomic analysis

For quantitative analysis of fatty acids and cholesterol, sciatic nerves were lysed and extracted in methanol/acetonitrile (MeOH:ACN) 1:1 v/v after addition of internal standards (IS: heneicosanoic acid, C21:0 for saturated fatty acids, ^13^C_18_-labeled linoleic acid C18:2 for unsaturated fatty acid and 5α-cholestane for cholesterol, Sigma-Aldrich). Then, the pellets were removed and used for total protein quantification by Bradford method. Fractions for the quantitative analysis of free cholesterol were first derivatized with a mixture of bis-trimethylsilyltrifluoroacetamide (BSTFA):pyridine (4:1 v/v) for 30 minutes at 60°C, and then injected into a gas chromatograph-mass spectrometer (GC-MS, Varian Saturn 2100). The MS was operated in the electron impact (EI) ionization mode. GC-MS analyses were performed as follows: 1μl sample was injected in splitless mode (inlet was kept at 270°C with the helium flow at 1.0 ml/min) at the initial 180°C. The oven was first kept at 180°C for 1 min, ramped at 50°C/min to 240°C, then at 5°C/min to 300°C for 6 min. The ions used for the quantification of cholesterol were at m/z 368 for cholesterol and m/z 357 for 5α-cholestane, the IS. The selection of ions for selective ion monitoring (SIM) analysis was based on mass spectra of pure standards and the quantification was based on calibration curves freshly prepared using a fixed concentration of the IS, and different concentrations of cholesterol, in a range from 0 to 10 μg/μl.

Total fatty acids were obtained from samples by acid hydrolysis. Briefly, chloroform/MeOH 1:1 v/v. and 1M HCl:MeOH (1:1, v/v) was added to the total lipid extracts and shook for 2h. After, chloroform:water (1:1 v/v) was added, and the lower organic phase was collected, transferred into tubes and dried under nitrogen flow. The residue was resuspended in 1 ml of MeOH (Cermenati, Abbiati et al., 2012). Fatty acid quantification was performed on an API-4000 triple quadrupole mass spectrometer (AB SCIEX) coupled with a HPLC system (Agilent) and CTC PAL HTS autosampler (PAL System)._The liquid chromatography mobile phases were (A) 10 mM isopropyl-ethyl-amine, 15 mM acetic acid in water:MeOH 97:3 (B) MeOH. The gradient (flow rate 0.5 ml/min) was as follows: T_0_: 20% A, T_20_: 1% A, T_25_: 1% A, T_25.1_: 20% A, T_30_: 20% A. The Hypersil GOLD C8 column (100mm × 3 mm, 3 μm) was maintained at 40°C. The injection volume was 10 μl and the injector needle was washed with MeOH/ACN 1:1 (v/v). Peaks off the liquid chromatography-tandem mass spectrometry (LC-MS/MS) were evaluated using a Dell workstation by means of the software Analyst 1.6.2. The mass spectrometer was operated in negative ion mode with the ESI source using nitrogen as sheath, auxiliary and sweeps gas, respectively. The mass spectrometer was operated in SIM/SIM based on mass spectra of pure standards and the quantification was achieved using calibration curves freshly prepared containing fixed concentration of the IS and different concentrations of the analyzed fatty acids (Cermenati, Audano et al., 2015)

### Bioavailability assay

3 WT and 3 S63del mice were treated with daily i.p. injection of BMS (48mg/Kg) for 10 days, starting at P15. Blood was collected in tubes with lithium heparin from three animals, centrifuged at 2,500 rpm at 4°C to isolate plasma. Sciatic nerves were collected, weighed and frozen with liquid nitrogen. To test BMS bioavailability samples were analyzed by liquid chromatography-tandem mass spectrometry with calibration standards (1 to 2,000 ng/mL) prepared from blank tissue homogenate (Eurofins ADME Bioanalyses, France)

### Statistical analysis

No statistical methods were used to predetermine sample sized, but our sample size are similar to those generally used in the field. Graphs and data were analyzed using GraphPad Prism Software. Data show the mean ± standard error of mean (SEM). One-way ANOVA with post hoc analysis (when specified in the text) was used. Significance levels were marked on figures as follows: P values (P), *P ≤ 0.05, **P ≤ 0.01, ***P ≤ 0.001.

## RESULTS

### Over-expression of Nrg1 type-III ameliorates electrophysiological and morphological parameters in a CMT1B mouse model

In many CMT neuropathies, the disease mechanism includes toxic gain of function (Scherer & Wrabetz, 2008). For example in S63del mice, a model of CMT1B that manifests developmental hypomyelination followed by progressive demyelination, the misfolded P0S63del protein accumulates in the ER and activates an unfolded protein response (UPR) (Pennuto et al., 2008, Wrabetz et al., 2006). To test whether augmenting the key pro-myelinating axonal signal Nrg1 type-III (TIII) (Taveggia et al., 2005) could overcome the intoxicated process and ameliorate hypomyelination, demyelination and remyelination, we crossed S63del mice with transgenic mice that over-express a HA-Nrg1TIII (HANI) fusion protein (Caroni, 1997, Velanac et al., 2012). Western blot (WB) analysis on spinal cord lysates, probed with anti HA antibody, confirmed the expression of the HANI transgene in HANI/+ and HANI/+//S63del mice (Supplementary Fig. 1A).

S63del mice manifest impaired motor capacity, reduced NCV and an increase FWL, similar to the alterations found in patients (Miller, Patzko et al., 2012, Wrabetz et al., 2006). Neurophysiological analysis on 6-month-old mice showed that overexpressing Nrg1TIII (HANI/+//S63del) in S63del mice resulted in a significant increase of NCV, that returned closer to WT values as compared to neuropathic mice (Fig. 1A and 1D). Similarly, the FWL was decreased to almost WT level in HANI/+//S63del (Fig. 1B and 1C). No significant modifications were observed in either cMAP amplitude or distal latency (Fig. 1D). These results indicate that genetic overexpression of Nrg1TIII leads to a significant amelioration of neurophysiological parameters in a mouse model of CMT1B.

**Figure 1.**
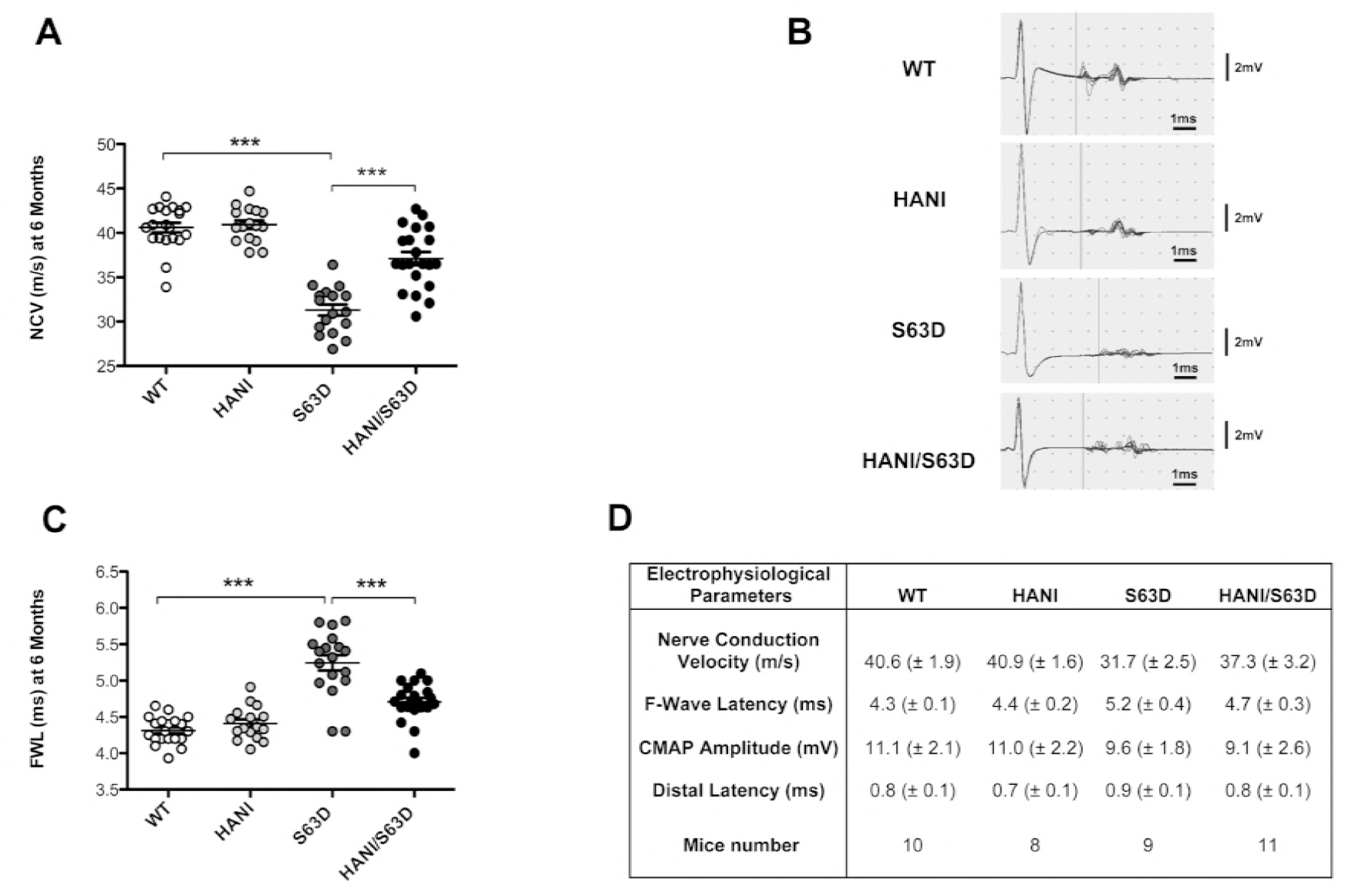
Over-expression of Nrg1 type III ameliorates neurophysiological parameters in P0S63del-CMT1B mice. Analyses were performed on 6 months old mice. **(A)** NCV is reduced in S63del mice as compared to WT and is rescued by Nrg1TIII overexpression. **(B)** Representative original recording indicating the onset of the F-wave (vertical bar). **(C)** F-wave latency is increased in S63del mice as compared to WT and shows a significant amelioration in HANI/+//S63del as compared to S63del mice. **(D)** CMAP Amplitude and Distal Latency do not show changes among the genotypes. Data represent mean ± SEM. Number of animals per genotype is reported in **D**. *** P value < 0.001 by one-way Anova with Bonferroni’s multiple comparison test.

To corroborate the electrophysiological findings, we performed morphological examinations of sciatic nerves starting at postnatal day 28 (P28). Transverse sections showed whole nerve hypertrophy restricted to peripheral nerves overexpressing the HANI transgene as compared to controls (Supplementary Fig. 1B). As expected, electron microscopy (EM) showed hypomyelination in S63del nerves as compared to wild type (Wrabetz et al., 2006), whereas HANI/+ nerves were clearly hypermyelinated (Fig. 2A) (Michailov et al., 2004, Velanac et al., 2012). Remarkably, myelin thickness was increased in HANI/+//S63del nerves as compared to S63del (Fig. 2A). Morphometric analysis confirmed these observations: g-ratios (axon diameter/fiber diameter) were decreased in HANI and HANI/+//S63del as compared to WT and S63del respectively (WT 0.65±0.006, HANI 0.54±0.012, S63del 0.69±0.005, HANI/+//S63del 0.62±0.002) (Fig. 2B and 2C). The increase in myelin thickness in HANI/+//S63del was predominant in the small to medium caliber axons (from 1 to 3.9μm) whereas those with larger diameter (more than 4μm) remained hypomyelinated with a g-ratio similar to S63del nerves (Supplementary Fig. 1C and 1D).

**Figure 2.**
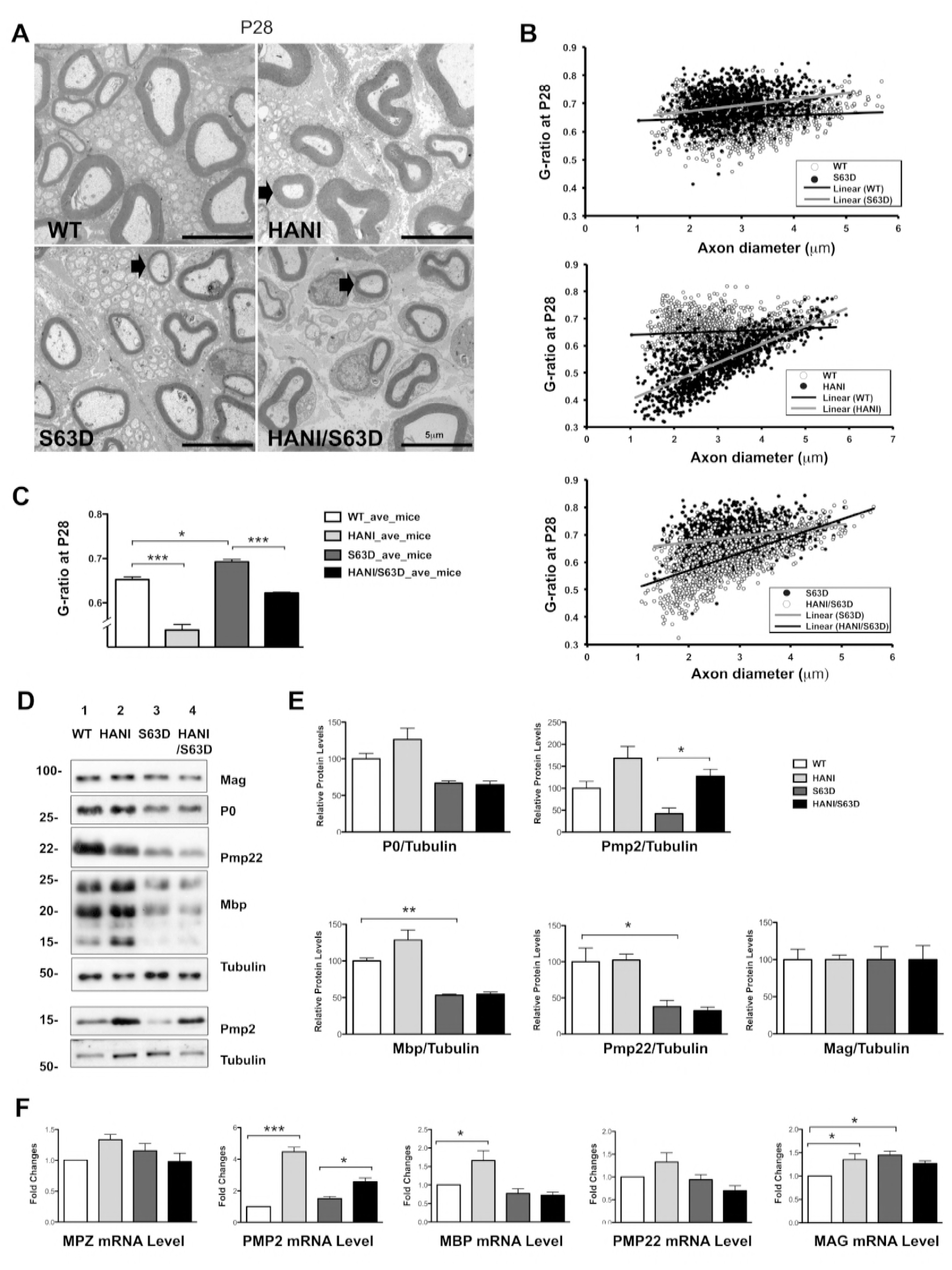
Over-expression of Nrg1 Type III leads to thicker myelin without increase in the expression of most myelin proteins. **(A)** Electron microscopy of P28 sciatic nerves. HANI mice show thicker myelin (arrow) as compared to WT. Hypomyelination in S63del is rescued in small and medium caliber axons (arrow) in HANI/+//S63del nerves (size bar 5μm). **(B)** G-ratio analysis performed on sciatic nerve cross sections at P28 and **(C)** relative quantification (average g-ratio values: WT 0.64 ± 0.006; HANI/+ 0.53 ± 0.012; S63del 0.69 ± 0.005; S63del//HANI/+ 0.62 ± 0.002). 8-10 microscopic fields per mouse were analyzed. 3-5 mice per genotype were used. **(D)** Western analysis on sciatic nerve lysates at P28 show no significant increase in myelin proteins in mice overexpressing Nrg1TIII, except for PMP2. Tubulin was used as loading control. One representative blot of three is shown **(E)** Protein levels as measured by densitometric analysis. **(F)** mRNA expression levels of myelin proteins measured via qRT-PCR. PMP2 mRNA is increased in mice overexpressing Nrg1TIII; MBP shows a small increase in HANI vs WT. *P < 0.05, **P < 0.01, ***P < 0.001 by one-way Anova with Bonferroni’s multiple comparison test; data represent the mean ± SEM. Each experiment was repeated three times on different pools of three nerves per genotype.

To evaluate whether Nrg1TIII overexpression was affecting myelin maintenance, we performed both morphological and morphometric analysis on sciatic nerves at 6-months: thicker myelin was still a feature of the HANI/+ mice (Supplementary Fig. 2A) and rescue of hypomyelination was again found in small and medium caliber axons in HANI/+//S63del mice (Supplementary Fig. 2B and 2C). Since CMT1B is characterized by progressive demyelination (Wrabetz et al., 2006) we also evaluated the number of demyelinated fibers and onion bulbs at 6-months of age, but we did not find significant differences in HANI/+//S63del as compared to S63del nerves (Supplementary Fig. 3C). This suggests that increased Nrg1TIII expression partially corrects hypomyelination without rescuing demyelination. All the aforementioned observations were confirmed at 12-months of age, indicating that, even at later stages, Nrg1TIII overexpression does not negatively affect myelin maintenance (Supplementary Fig. 3D).

### Overexpression of Nrg1TIII may act through a Krox20 independent pathway

During peripheral nerve development the binding of Nrg1TIII to ErbB tyrosine kinase receptors (ErbB2/3 heterodimers) on the Schwann cell surface, triggers the activation of intracellular signaling pathways, such as PI3K/p-Akt and Erk1/2-MAPK, that lead to myelin protein and lipid gene expression, mostly acting through the transcription factor Krox20 (Leblanc et al., 2005, Parkinson et al., 2004). It would be expected that the increase in myelin thickness due to Nrg1TIII overexpression should be accompanied by a concomitant increase in myelin proteins or lipids. We therefore performed both gene expression and biochemical analysis on P28 sciatic nerves extracts for the major proteins of compact and non-compact myelin. Surprisingly, Western blot (WB) as well as mRNA analysis showed only a small increase of P0, Mbp, Pmp22 and Mag expression in HANI/+ as compared to WT, whereas no differences were found in HANI/+//S63del as compared to S63del mice (Fig. 2D-F). Interestingly, only peripheral myelin protein 2 (Pmp2, P2 or Fabp8), a protein with lipid binding activity in peripheral nerve myelin (Chrast, Saher et al., 2011, Uyemura, Yoshimura et al., 1984, Zenker, Stettner et al., 2014), showed a strong increase of gene and protein expression in HANI/+ and HANI/+//S63del mice as compared to controls (Fig. 2D-F). Similar results were found in sciatic nerves lysates at 6 months of age, where we could detect only a slight increase of P0 and Pmp22 protein expression in S63del mice overexpressing the HANI transgene (Supplementary Fig. 4). Overall, our data indicated that despite Nrg1TIII overexpression there is only a minor effect on myelin mRNA and proteins. Therefore, we checked for the activation of the pathways downstream Nrg1TIII. The HANI transgene is highly expressed in spinal cord extracts (Supplementary Fig. 1A), but the HA-tag is barely detectable in nerves (Velanac et al., 2012) (data not shown). However, by probing P28 sciatic nerve extracts with an antibody against the C-terminus of Nrg1 (all isoforms), we could detect, especially in the S63del background, an increase of the cleaved active isoform of Nrg1 in mice expressing the transgene (Fig. 3B). As expected the levels of phosphorylation of both Akt^(Ser473)^ and Erk1/2 were significantly increased in HANI/+ and HANI/+//S63del nerves as compared to controls, suggesting that HA-tagged Nrg1TIII was correctly activating the PI3K and MAPK signaling pathways (Fig. 3B). Paradoxically however, we did not find any increase of Krox20 mRNA and protein expression (Fig. 3B), which could explain why myelin genes and proteins are not significantly affected in sciatic nerves overexpressing Nrg1TIII. This may suggest that Nrg1TIII rescues hypomyelination in S63del nerves through a Krox20-independent mechanism. Finally, no significant differences in phospho-Akt^(Ser473)^, phospho-Erk1/2 and Krox20 levels were detected at 6 months of age (Supplementary Fig. 3A and 3B and data not shown), suggesting that Nrg1TIII signalling gets attenuated with time.

**Figure 3.**
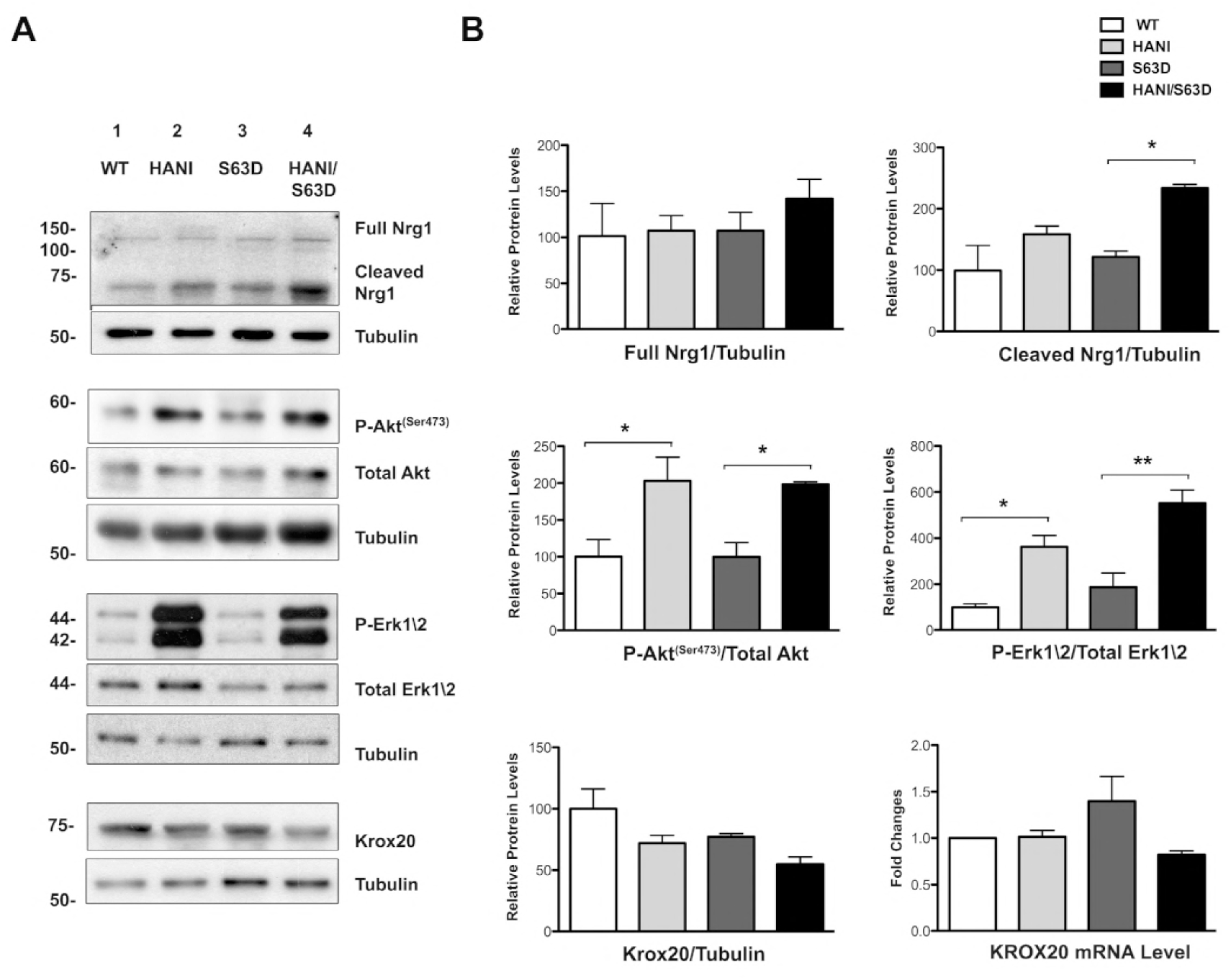
The Akt and Erk1/2 signaling pathways are upregulated by Nrg1 TIII overexpression. **(A)** Western blot analysis of P28 sciatic nerves lysates. No differences in the endogenous full-length form of Ngr1 (140KDa) were detected amongst the four genotypes, whereas the cleaved (65-70KDa) active form was increased in particular in HANI/+//S63del as compared to S63del. In HANI/+ and HANI/+//S63del mice both Akt^(Ser473)^ and Erk1/2 phosphorylation are increased as compared to S63del and WT mice, respectively. Krox20 protein expression is slightly, but not significantly decreased in HANI/+//S63del as compared to S63del mice. No differences were detected in Krox20 mRNA expression by qRT-PCR (B, lower right graph). Tubulin was used as loading control. One representative experiment of three is shown. **(B)** Densitometric quantification of full-length and cleaved Nrg1 forms, p-Akt^(ser473)^, p-Erk1/2 and Krox20 protein levels. * P < 0.05 and ** P < 0.01 by one-way Anova with Bonferroni’s multiple comparison test.

### Nrg1 type III overexpression increases myelin lipid content

One possible explanation for the increase in myelin thickness without corresponding increase in myelin proteins is an alteration in myelin packing; we thus checked myelin periodicity in sciatic nerves at P30. No gross alteration of the myelin sheath periodicity was found among all four genotypes (Supplementary Fig. 5), suggesting that myelin is normally compacted. These observations could therefore be predictive of an altered protein density in the myelin sheath. Indeed, immuno-EM analysis (IEM) revealed a trend towards a reduction of P0 density in the myelin of mice overexpressing Nrg1TIII, particularly in the S63del background (Fig. 4A and 4B).

**Figure 4.**
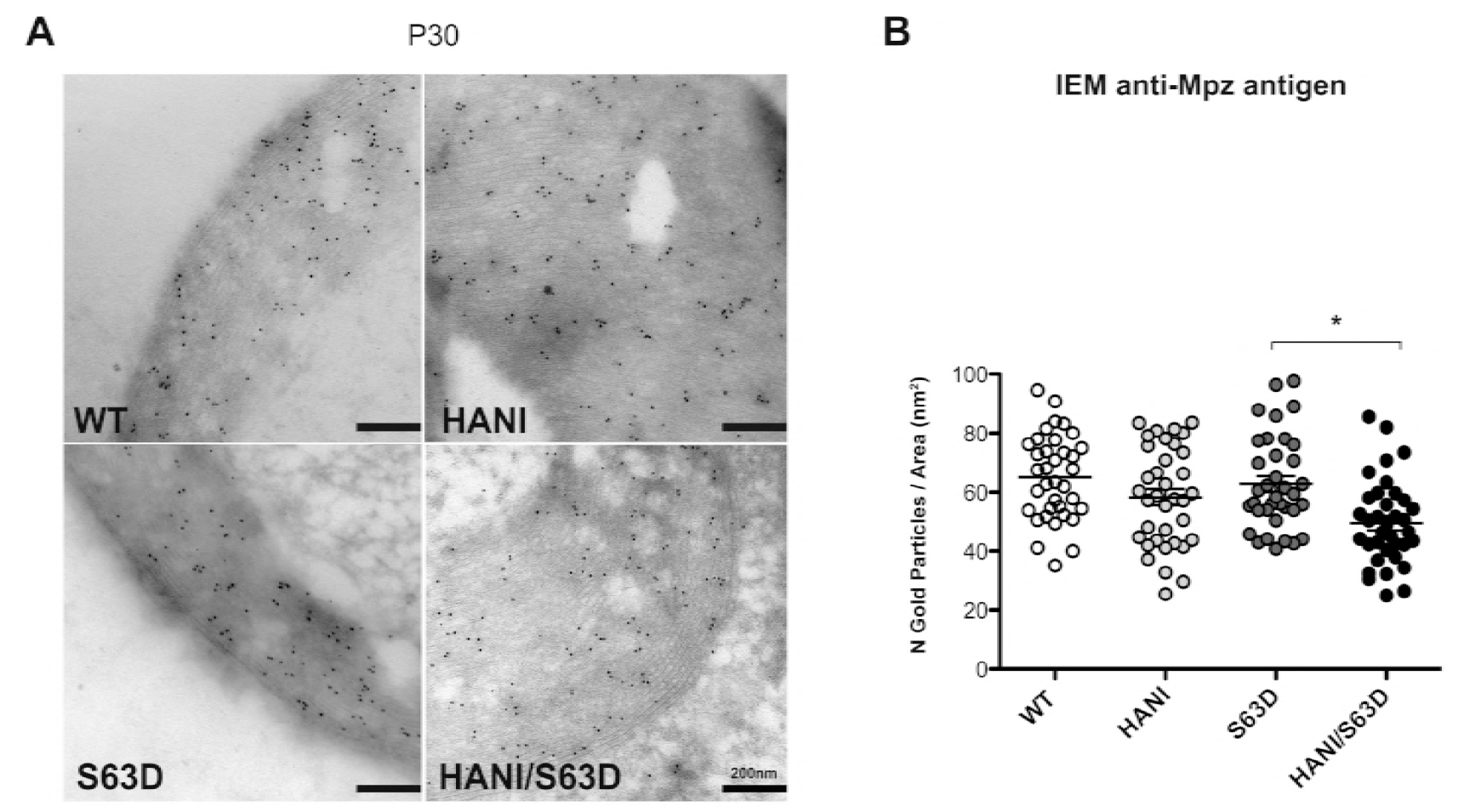
P0 protein density is reduced in mice expressing the HANI transgene. **(A)** IEM for P0 in transverse sections of P30 sciatic nerves. **(B)** Quantification of the number of gold granules per myelin area shows a trend towards the reduction of P0 density in HANI/+ and HANI/+//S63del myelin as compared to WT and S63del, respectively. Twelve sciatic nerve images per genotype were counted from three mice per genotype. * P value = 0.03 by Student’s T-test; error bars represent SEM.

In contrast to most biological membranes, myelin is characterized by a higher lipids to proteins ratio (~75:25) (Greenfield, Brostoff et al., 1973, Siegel, 1999). Thicker myelin, with reduced protein density could imply a relative increase of the total lipid content. To test this hypothesis, we performed a lipidomic analysis on P28 sciatic nerves. We detected increased levels of free cholesterol (typical of the myelin sheath) in mice overexpressing the HANI transgene as compared to WT, whereas no significant differences were found between S63del and HANI/+//S63del mice (Fig. 5A). Moreover, most saturated (Palmitic acid C16:0; Stearic acid C18:0; Behenic C22:0 and Lignoceric C24:0) and unsaturated (Oleic acid C18:1; Linoleic acid C18:2; Erucic acid C22:1 and Nervonic acid C24:1) fatty acids showed increased levels in mice harboring the HANI transgene (Fig. 5B). Lipids are important determinants of myelin structure and function, and proper lipid stoichiometry can be indicative of myelin integrity. We calculated the desaturation index (ratio 18:0/18:1) and the membrane fluidity index (ratio 18:1/18:2) (Chrast et al., 2011) that showed no alteration, in accordance with a thicker, but still properly compact myelin. Taken together these data suggest that overexpression of Nrg1TIII determines a thickening of the myelin sheath that appears to be more dependent on an increase in lipids rather than proteins content.

**Figure 5.**
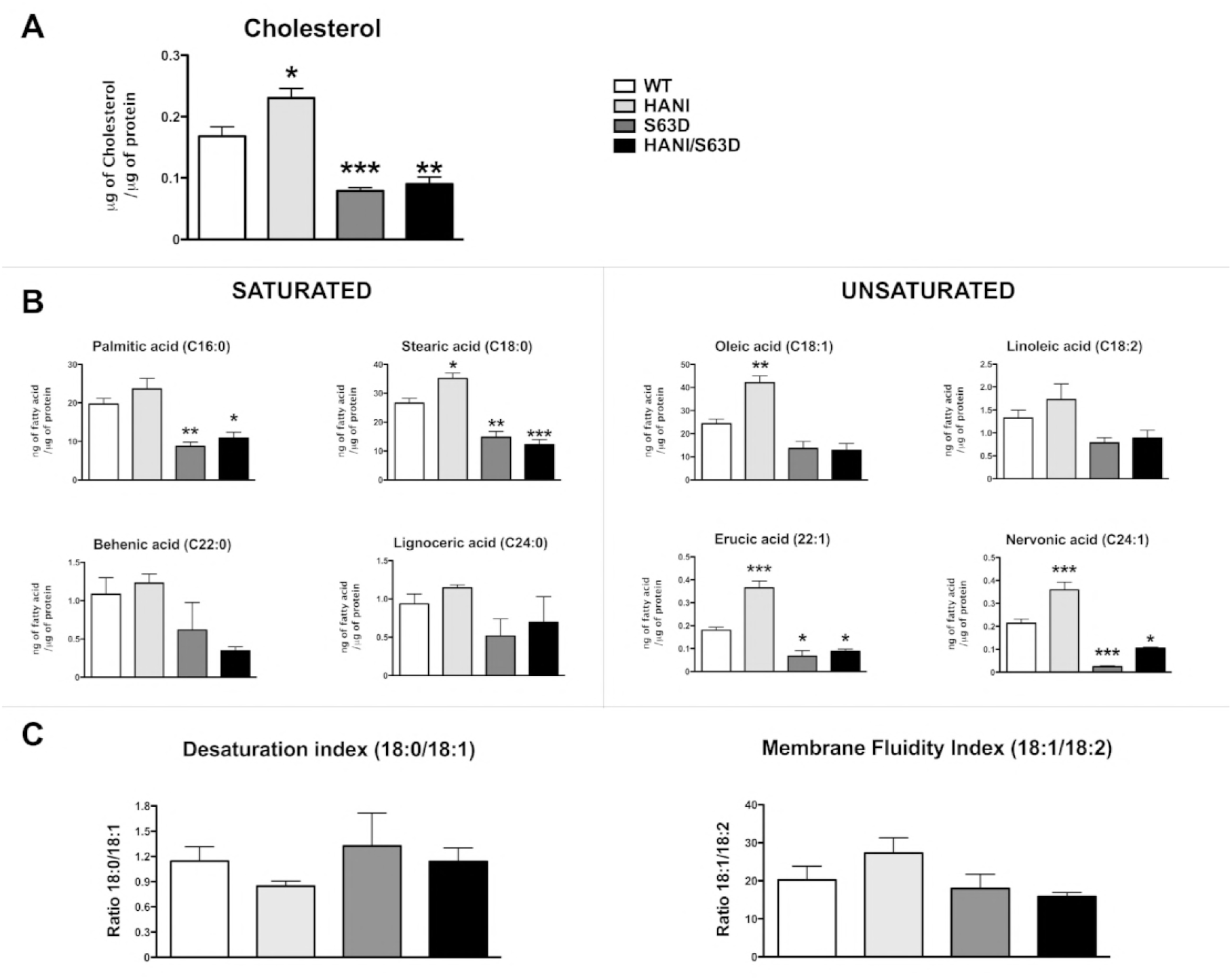
Nrg1TIII overexpression increases cholesterol and fatty acids levels. Lipidomic analysis was performed on P28 sciatic nerves. **(A)** The level of free cholesterol increased in HANI as compared to WT, whereas no differences were detected between S63del and HANI/+//S63del nerves. **(B)** Levels of total saturated (Palmitic acid C16:0; Stearic acid C18:0; Behenic C22:0 and Lignoceric C24:0) and total unsaturated (Oleic acid C18:1; Linoleic acid C18:2; Erucic acid C22:1 and Nervonic acid C24:1) fatty acids and **(C)** the relative Desaturation Index (ratio 18:0/18:1) and Membrane Fluidity Index (ratio 18:1/18:2). Data are expressed as μg of cholesterol or ng of fatty acid normalized to μg of total proteins. * P < 0.05, ** P < 0.01, *** P < 0.001 *vs* WT by one-way Anova with Bonferroni’s multiple comparison test. Five sciatic nerves from different animals per genotype were analyzed.

### Nrg1TIII overexpression does not alter the stress response in S63del Schwann cells

P0S63del is a misfolded protein that fails to be incorporated into myelin and is retained in the ER where it activates a dose-dependent UPR (Pennuto et al., 2008, Wrabetz et al., 2006). In our model, P0S63del is expressed by an *MPZ*-based transgene that faithfully parallels WT P0 expression (Feltri, D’Antonio et al., 1999). It was therefore conceivable that overexpressing Nrg1TIII in S63del mice could increase P0 mutant protein expression, further intensifying the toxic gain of function and potentially worsening the phenotype. However, we observed an amelioration of the neuropathic phenotype coupled to a lack of increase in myelin genes and proteins expression. Importantly, an allelic discrimination assay showed that the ratio between WT and mutant P0 mRNA remained unaltered in S63del and HANI/+//S63del nerves (Supplementary Fig. 6D) strongly suggesting that, just like the other myelin proteins, P0S63del expression was not significantly affected by Nrg1TIII overexpression. To further assess this, we analyzed the expression of UPR markers in HANI/+//S63del and littermate control mice at P28 and 6 months (Fig. 6 and Supplementary Fig. 6). As previously shown, BiP, Grp94 and P-eIF2alpha proteins, as well as BiP, Chop and Xbp1s mRNAs were strongly increased in S63del nerves (D’Antonio et al., 2013, Pennuto et al., 2008, Wrabetz et al., 2006). However, we did not detect any change in the expression of these UPR markers in HANI/+//S63del indicating that after Nrg1TIII overexpression the UPR was still active at levels comparable to those of S63del nerves. Thus, Nrg1TIII overexpression ameliorates the phenotype of a CMT1B mouse model without further intensifying the toxic gain of function mechanism.

**Figure 6.**
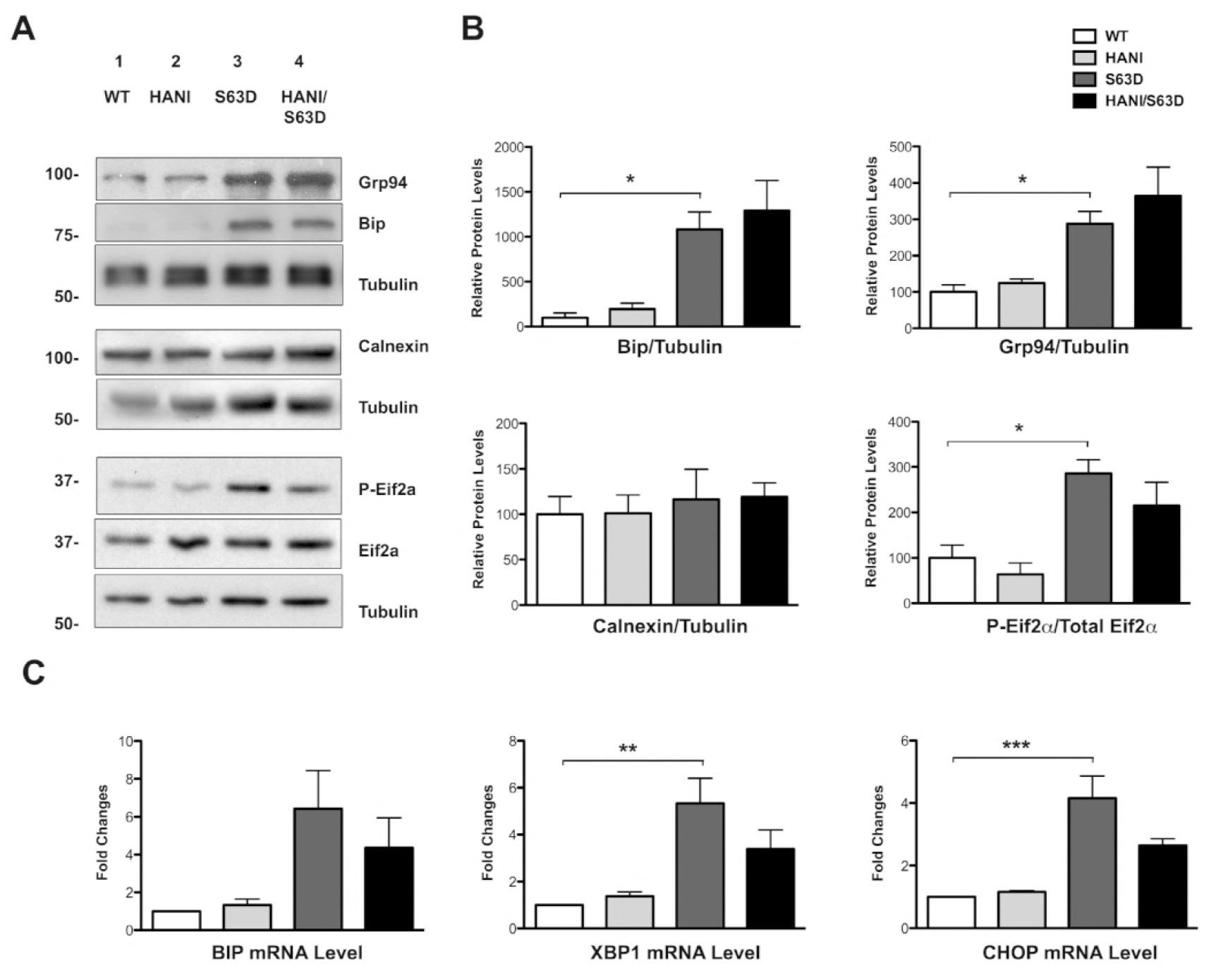
ER stress levels do not increase in S63del/HANI/+ mice. **(A)** Western blot analysis of P28 sciatic nerves lysates shows activation of the ER stress markers Bip/Grp78, Grp94 and P-eIF2-alpha in S63del mice as compared to WT. No further increase was detected in the expression levels of these markers between S63del and HANI/+//S63del mice. Tubulin was used as loading control. One representative experiment of three is shown. **(B)** Densitometric quantification of Bip, Grp94, Calnexin and p-eIF2-alpha relative protein levels. **(C)** qRT-PCR for BiP, CHOP, and spliced XBP-1 mRNA in P28 sciatic nerves. All the stress makers are increased in S63del as compared to WT, but we detected no differences in their levels of expression between S63del and HANI/+//S63del nerves. Each experiment was repeated three times on different pools of three nerves per genotype. * P < 0.05, ** P value < 0.01 and *** P < 0.001 by one-way Anova with Bonferroni’s multiple comparison test.

### Inhibition of TACE ameliorates myelination in neuropathic DRG explant cultures

Taken together our data suggest that the modulation of Nrg1TIII activity may represent a suitable approach to treat *MPZ*-related neuropathies. La Marca and colleagues have shown that the α-secretase TACE cleaves Nrg1TIII thus limiting the amount of functional Nrg1TIII expressed on the axonal surface and inhibiting myelination in early development (La Marca et al., 2011). TACE null mice are hypermyelinated, partially mimicking the phenotype observed in transgenic mice overexpressing Nrg1TIII. This, together with our data, predicts that inhibitors of TACE would activate Nrg1TIII and promote myelination. BMS-561392 is a highly selective TACE inhibitor, already employed in phase I and II clinical trials, as a treatment for rheumatoid arthritis (Grootveld & McDermott, 2003, Moss, Sklair-Tavron et al., 2008). In a preliminary set of experiments, we treated P15 WT and S63del mice with daily i.p. injections of BMS-561392 for 10 days. Unfortunately, bioavailability analysis revealed almost undetectable levels of BMS in both WT and S63del sciatic nerves suggesting that this drug does not efficiently cross the blood nerve barrier (BNB) (Supplementary 7B). Thus, to test the potential of BMS-561392 in promoting myelination, we used organotypic explant cultures of myelinating dorsal root ganglia (DRGs), a reliable model of *ex-vivo* myelination. We treated WT and S63del DRGs with 1μM BMS-561392 for 2-weeks along with controls (untreated and DMSO-treated DRGs) (Fig. 7A). Staining for myelin basic protein (MBP) clearly showed an increase in the number of myelinated segments in both S63del (Fig. 7C) and WT treated samples (Fig. 7B and Supplementary Fig. 7A), indicating that TACE inhibition ameliorates myelination, and suggesting that novel inhibitors with improved BNB permeability could represent promising therapeutic candidates for hypomyelinating CMTs.

**Figure 7.**
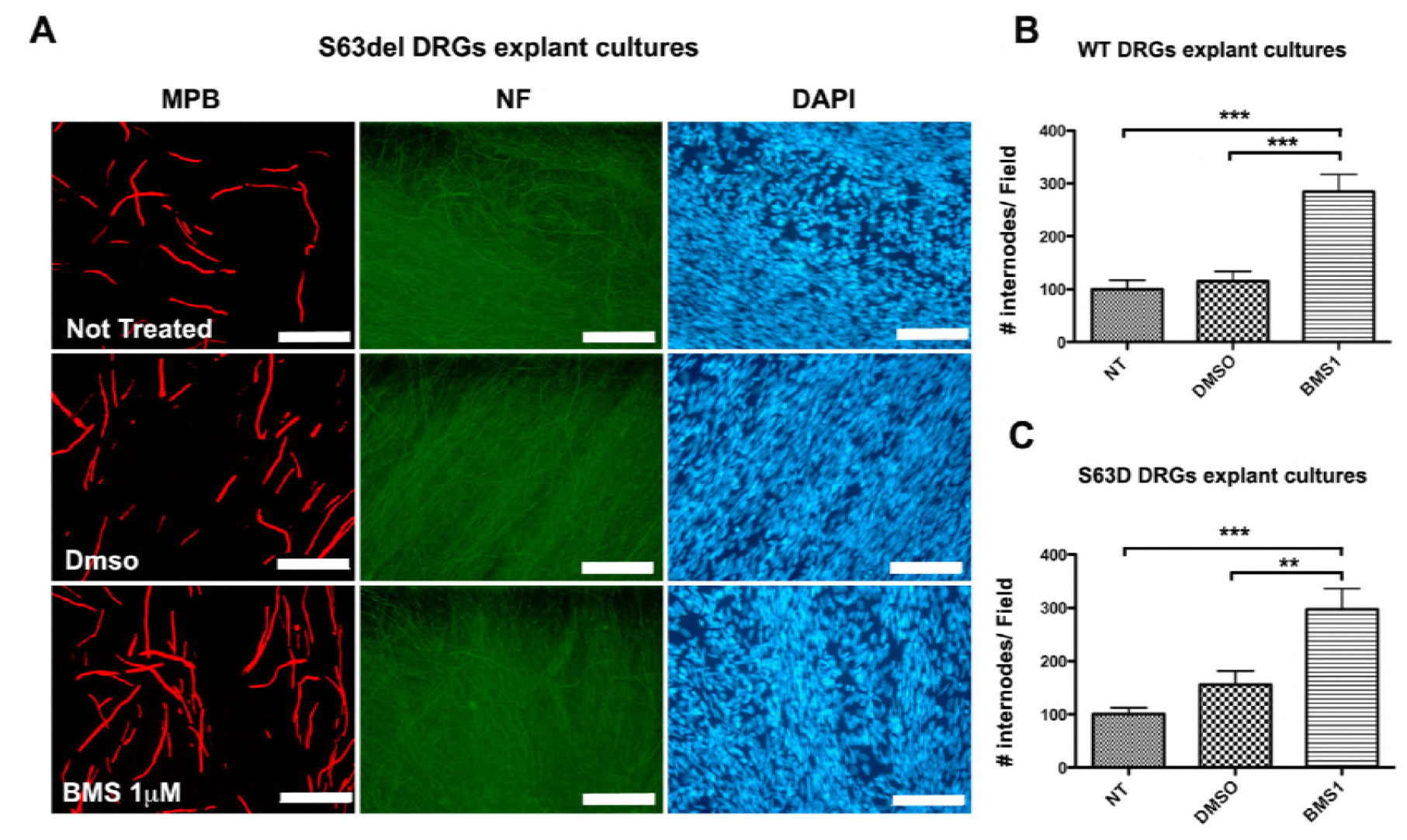
Pharmacological treatment with the TACE inhibitor BMS-561392 ameliorates myelination in DRGs explant cultures. Myelinating DRGs were treated with 1uM BMS (BMS1) for 2 weeks. As controls, not treated and DMSO-treated DRGs were analyzed. **(A)** Immunostaining for MBP showing the increase in the number of MBP-positive segments in S63del treated samples (size bar 100μm). Quantification of internode number in **(B)** WT (see also Supplementary Fig. 7A) and **(C)** S63del DRGs. Note how treatment with BMS increases the number of internodes in both the genotypes. Ten to fifteen DRGs per condition from three independent dissections were analyzed. **P < 0.01 and *** P < 0.001 by one-way Anova with Bonferroni’s multiple comparison test; error bars represent SEM.

## DISCUSSION

The analysis of the molecular mechanism underlying CMTs has revealed an extensive variety in pathogenesis (Baets, De Jonghe et al., 2014, Jerath & Shy, 2015, Wrabetz et al., 2006) so that the development of a specific therapy for each subtype would be difficult to achieve. The identification of common approaches for the treatment of hereditary neuropathies, independently of their pathogenetic mechanism, is therefore highly desirable.

As an essential regulatory signal driving myelination in the PNS (Michailov et al., 2004, Taveggia et al., 2005), Nrg1TIII is an excellent candidate for the development of general therapeutic approaches for CMTs. Indeed, recent work has shown that the genetic or pharmacological reduction of Nrg1TIII signaling rescues the phenotype in mouse models characterized by focal hypermyelination (Bolino, Piguet et al., 2016).

Relying on the same concept we hypothesized that increasing Nrg1TIII could ameliorate demyelinating neuropathies characterized by reduced levels of myelination. However, whereas the increase of Nrg1 signaling is expected to rescue neuropathies where loss-of-function is the prevalent mechanism (see accompanying paper from Belin et al.), in the presence of a toxic myelin protein it could potentially have detrimental effects. Here, we showed that the genetic overexpression of Nrg1TIII ameliorates the neuropathic phenotype of a preclinical mouse model of CMT1B characterized by P0 misfolding and UPR activation (Pennuto et al., 2008, Wrabetz et al., 2006) without aggravating the toxic gain-of-function. In this context, Nrg1TIII appears to act through a Krox20-independent mechanism, which determines a readjustment in myelin protein:lipid stoichiometry. Finally, we provide proof of principle that pharmacological enhancement of Nrg1 signaling may represent an appealing therapeutic approach.

### Nrg1TIII overexpression activates Krox20-independent signaling pathways

PI3K/Akt and MAPK/Erk1/2 pathways are known downstream effectors of Nrg1 and thought to trigger Krox20-dependent myelin and lipids genes activation (Leblanc et al., 2005, Parkinson et al., 2004) For instance, the upregulation of Akt phosphorylation has been directly linked to an increased activation of Nrg1TIII, both *in vivo* and *in vitro* (La Marca et al., 2011, Maurel & Salzer, 2000, Ogata, Iijima et al., 2004). However, following HANI transgene expression, the activation of both Akt^(Ser473)^ and Erk1/2 was uncoupled from Krox20 and myelin genes levels. Intriguingly, recent work has revealed a Krox20-independent signaling pathway in a mouse model overexpressing a constitutively active Akt1 isoform (MyrAkt) in Schwann cells. The regulatory mechanism appears to involve mTORc1 in regulating myelin thickness in the PNS, without affecting Egr2/Krox20 transcription levels (Domenech-Estevez, Baloui et al., 2016). Interestingly, a mild but sustained constitutive expression of the protein kinase Mek1 in Schwann cells (Mek1DD mouse), leads to a progressive and redundant hypermyelination in peripheral nerves. In these mice, the enhancement of Erk1/2 phosphorylation, but not Akt, induces robust protein synthesis without significantly affecting myelin mRNA levels. This effect is partially mediated by an mTORc1-dependent regulatory mechanism (Sheean, McShane et al., 2014). These observations corroborate the idea that the exogenous overexpression of Nrg1TIII could enhance myelin growth in sciatic nerves through either the PI3K/Akt^(Ser473)^ or the MAPK/Erk1/2 pathway, independently from Krox20 and myelin genes levels. However, in our context we did not observe any significant alteration downstream of the mTORc1 signaling pathway (data not shown), thus ruling out that the HANI transgene overexpression may act through mTOR to enhance protein translation, and suggesting a different mechanism, possibly acting on lipids (see below).

The uncoupling of Krox20 from upstream Nrg1TIII signaling may paradoxically underlie the rescue in the S63del/HANI nerves. In fact, in S63del mice, the UPR and the resulting neuropathic phenotype are remarkably dose dependent. Mice with higher levels of expression of P0S63del (S63del-H mice, 200% over-expression) have higher stress levels and a very severe phenotype as compared to the S63del mice used in this study (S63del-L, 60% over-expression) (Wrabetz et al., 2006). The fact that, like most myelin genes the expression levels of P0S63del were not directly affected, as confirmed by lack of increase in UPR markers, probably allowed Schwann cells to increase myelination without further intoxicating the system.

### The overexpression of Nrg1TIII might determine a shift in myelin proteins:lipids stoichiometry

The thicker myelin of HANI overexpressing mice had no alterations in periodicity and compaction, despite the reduction in major myelin proteins, as epitomized by the decrease of P0 protein density. This suggests a general change in myelin composition. Normally myelin is composed by 20% of proteins and by 80% of lipids such as free-cholesterol and fatty acids (Chrast et al., 2011, Garbay, Heape et al., 2000, Greenfield et al., 1973, Schmitt, Castelvetri et al., 2015). In the mice overexpressing Nrg1TIII, cholesterol along with saturated and unsaturated fatty acids was increased. Interestingly, the increase in lipids appeared more evident in the WT than in S63del background. This could depend on the fact that P0S63del expression leads to a general down regulation in cholesterol and lipid synthesis genes (D’Antonio et al., 2013), which may partially counteract the effects of Nrg1TIII. In agreement with our observations, recent work has identified the transcription factor Maf as a crucial player in regulating myelination and cholesterol biosynthesis downstream from Nrg1 (Kim, Wende et al., 2018).

Cholesterol is one of the most important regulators of lipid organization in the membrane and mice that lack cholesterol biosynthesis in glia cells show abnormal myelin structures (Saher, Quintes et al., 2009, Saher, Quintes et al., 2011). It is intriguing to notice that Pmp2, a member of the fatty acid binding protein family (FABPs) with a role in the binding and transport of lipids to the membrane (Chrast et al., 2011), was the only myelin protein strongly increased by Nrg1TIII overexpression. While Pmp2 does not appear to have a role in myelin structure, it may be important for myelin lipids homeostasis (Zenker et al., 2014). Thus, it is possible that high Pmp2 protein levels could enhance lipids transport and myelin compaction (Chrast et al., 2011, Majava, Polverini et al., 2010, Zenker et al., 2014). Interestingly similar observations were made in the Mek1DD mice (Sheean et al., 2014), which suggest that PMP2 is mostly regulated through the MAPK/ERK1/2 axis.

The increase in myelin thickness and the change in myelin composition, however, did not appear to affect demyelination. We have previously shown that demyelination in the S63del mice can be corrected by ablation of the UPR effectors Chop and Gadd34 (D’Antonio et al., 2013, Pennuto et al., 2008). The fact that the UPR is virtually unchanged by Nrg1TIII overexpression may explain why demyelination is not ameliorated.

The Nrg1TIII-driven reduction in g-ratio was more pronounced for axon with small/medium diameter. The reasons for this are not entirely clear, but it has been previously hypothesized that it may depend on the fact that an exponential increase in myelin synthesis, possibly not provided by the Thy-1.2 promoter, would be required to match the linear increase in axon diameter (Michailov et al., 2004). Still, the rescue of hypomyelination in this subset of axons was sufficient to significantly ameliorate important neurophysiological parameters, such as NCV and FWL. Unfortunately, we were not able to measure whether there was also an improvement in motor function. In fact the mice expressing the HANI transgene, even in a wild-type background, were incapable to perform well on Rotarod. This may depend on the expression of the HANI transgene in areas of the central nervous system such as cerebellum and hippocampus (Caroni, 1997), which may affect the equilibrium and suggests that expression of Nrg1TIII should be locally confined to the PNS. However, it should also be noted that recent works suggest that Nrg1 ameliorates cognitive function impairments and neuropathology in different Alzheimer’s disease (AD) mouse models, indicating that it may serve as potential candidate for the prevention and treatment of AD (Jiang, Chen et al., 2016, Ryu, Hong et al., 2016, Xu, de Winter et al., 2016).

### Neuregulin modulation as therapeutic target

Overall our data indicate that Nrg1TIII increase may partially overcome the myelin defects due to the expression of a toxic myelin protein. Notably, recent work has indicated that the genetic overexpression of the Nrg1TI, but not of the Nrg1TIII isoform, was effective in ameliorating the impaired nerve development defect in the EC61-CMT1A-CH mouse model (Fledrich et al., 2014). The reason of this discrepancy could rely on the fact that Fledrich and colleagues showed an imbalanced activity of the PI3k/Akt and the Mek/Erk1/2 signaling pathways in CMT1A models, that leads Schwann cells to acquire a persistent differentiation defect during early postnatal development, leading to dysmyelination (Fledrich et al., 2014). In contrast, in the S63del-CMT1B mouse, despite an increase of dedifferentiation markers (D’Antonio et al., 2013, Florio, Ferri et al., 2018), we never observed any imbalance between PI3K/Akt and Mek/Erk1/2 activation.

Nrg1TIII shedding by the α-secretase TACE/ADAM17 negatively regulates PNS myelination (La Marca et al., 2011). Whereas directly increasing Nrg1 may have detrimental side effects, the selective modulation of its secretase is an interesting therapeutic approach for the treatment of human neuropathies. Indeed, a recent study showed promising results in treating CMTs mouse models characterized by excessive redundant myelin thickness with Niaspan, a FDA-approved drug known to enhance TACE/ADAM17 activity (Bolino et al., 2016). Similarly, treatment of P0S63del DRG explant cultures with the TACE inhibitor BMS-561392 showed promising results in ameliorating myelination. Future goals will include the synthesis of specific chemical compounds able to cross the barrier and target the nerves.

### Conclusions

We provided proof of principle that modulating Nrg1 is a potential general therapeutic approach for hereditary neuropathies. The exogenous overexpression of Nrg1TIII ameliorated the neuropathic phenotype of a CMT1B mouse model characterized by accumulation of misfolded P0S63del and activation of the UPR, without exacerbating the toxic mechanism. We also showed that Nrg1TIII overexpression may act through a Krox20-independent pathway that affects myelin lipids synthesis rather than protein content. The identification of new small molecules that directly modulate Nrg1TIII activity will have relevant clinical implications to treat hypomyelinating CMTs.

## Acknowledgements

We thank Maria Carla Panzeri and the Alembic facility at the San Raffaele Scientific Institute for excellent technical support. This work was supported by grants from the National Institute of Health (LW R01 NS55256 and R56NS096104), the Charcot Marie Tooth Association (LW), Fondazione Telethon (GPP10007 to LW, MLF and CT, GGP14147 to MDA and GGP15012 to MDA and CT) the Italian Ministry of Health (GR-2011-02346791 to MDA) and the Umberto Veronesi Foundation (Fellowship Award 2014 to CS). MHS holds a Heisenberg Fellowship from the Deutsche Forschungsgemeinschaft (DFG) and acknowledges funding by a DFG research grant (SCHW741/4-1).

## SUPPLEMENTARY FIGURES LEGENDS

**Supplementary Figure 1.**
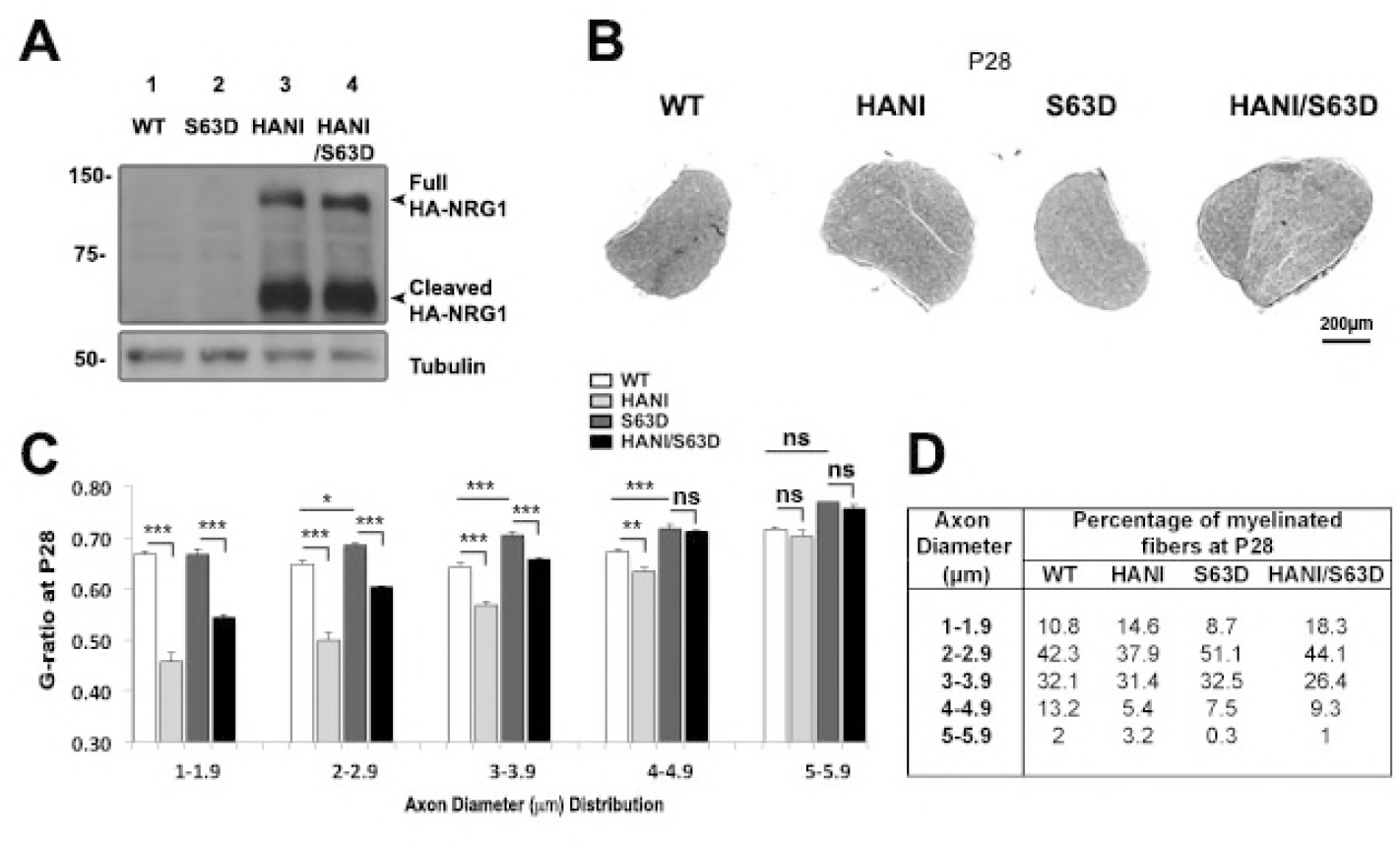
**(A)** Expression of Nrg1TIII-HA (HANI) transgene in mice at P28. Western analysis on P28 spinal cord lysates probed with the anti HA antibody. The bands corresponding to full length and cleaved Nrg1TIII are detected only in mice expressing the transgene. **(B)** Transverse semi-thin cross sections of sciatic nerves at P28 show hypertrophy in HANI/+ and HANI/+//S63del mice as compared to controls (size bar 200μm). **(C)** G-ratio plotted by axon diameter in sciatic nerves at P28 shows that the rescue of hypo-myelination is restricted to small and medium (from 1 to 3.9μm) caliber axons.* P < 0.05, ** P < 0.01, *** P < 0.001 and *ns* (not significant) by one-way Anova with Bonferroni’s multiple comparison test. **(D)** Percentage of myelinated fibers plotted versus axons diameter distribution at P28.

**Supplementary Figure 2.**
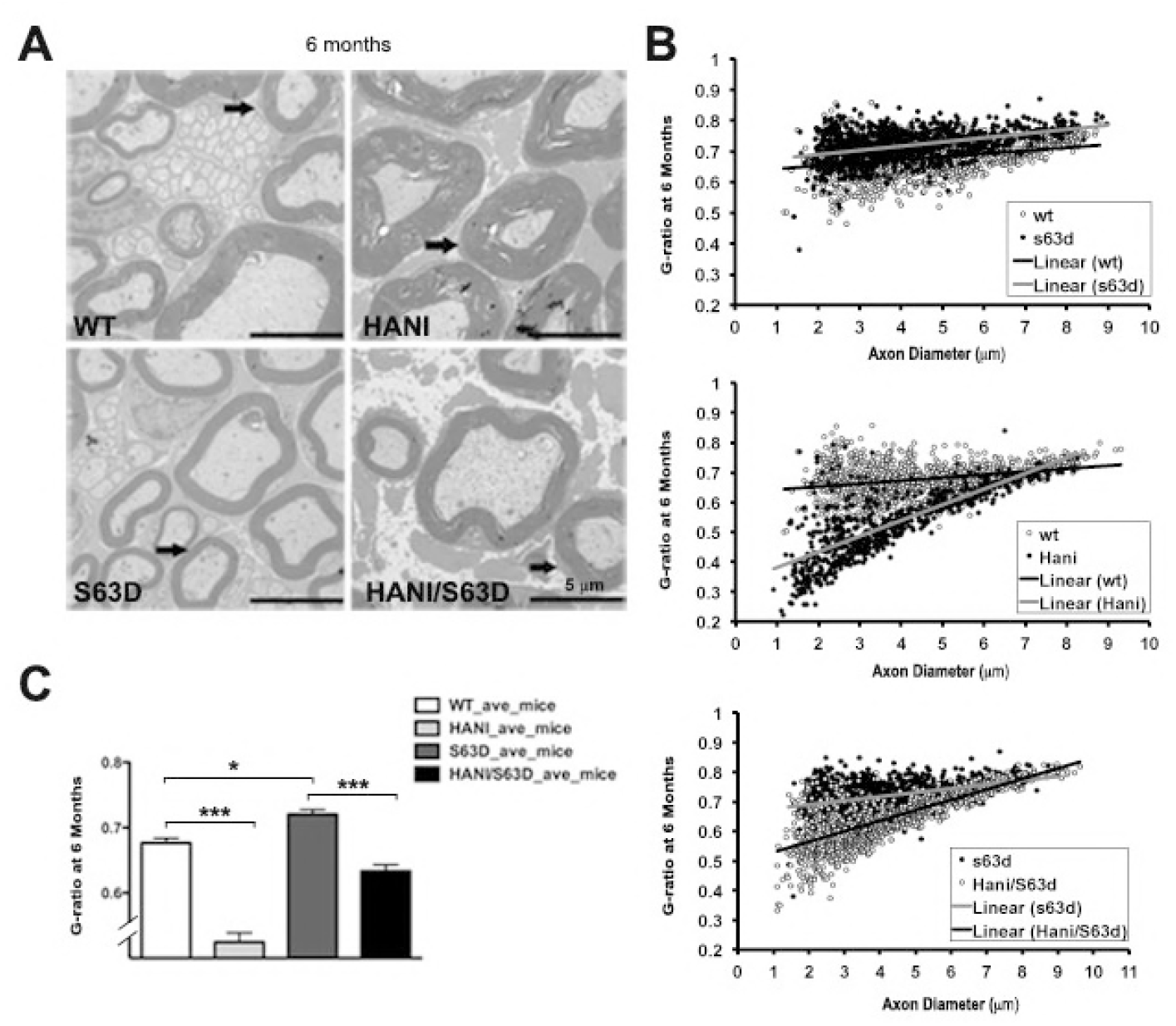
**(A)** Electron microscopy of sciatic nerve transverse sections at 6 months shows hypomyelination in S63del mice as compared to WT, whereas HANI mice show thicker myelin. Hypomyelination is still rescued in small and medium caliber axons in HANI/+//S63del mice as compared to S63del controls (size bar 5μm). **(B)** G-ratio analysis performed on sciatic nerve cross sections at 6 months and **(C)** average g-ratio values: WT 0.68 ± 0.007; HANI/+ 0.52 ± 0.014; S63del 0.72 ± 0.007; HANI/+//S63del 0.63 ± 0.01; data represent the mean ± SEM. At least three mice per genotype were analyzed. * P < 0.05, *** P < 0.001 by one-way Anova with Bonferroni’s multiple comparison test.

**Supplementary Figure 3.**
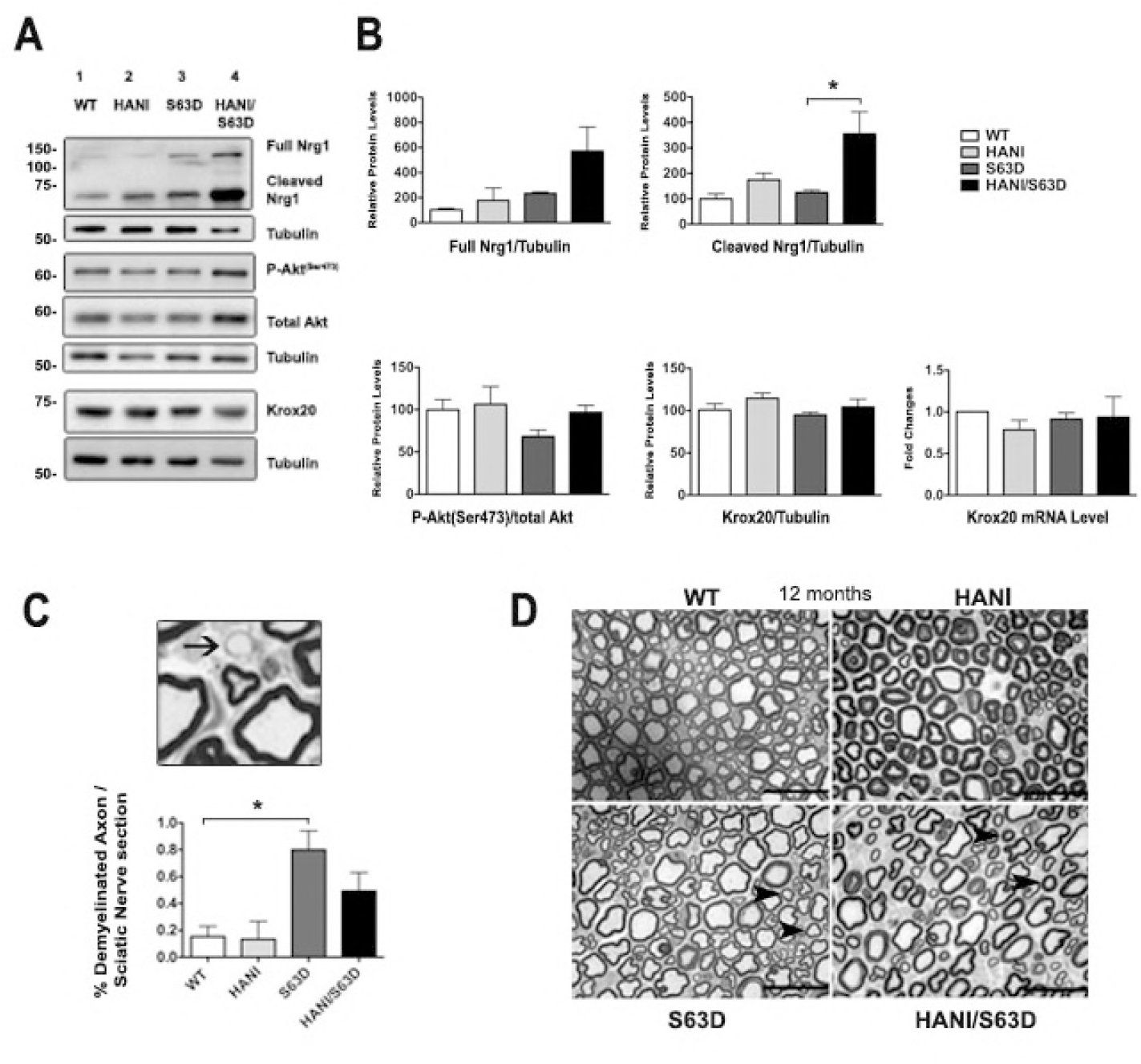
**(A)** There are no differences in full-length Nrg1 among the genotypes and a significant increase in the cleaved form of Nrg1 in HANI/+//S63del as compared to neuropathic mice in 6 months old sciatic nerves by Western blot analysis. No differences were detected in the levels of phospho-Akt^(Ser473)^ and for Krox20 protein expression. **(B)** Quantification of the full-length and cleaved Nrg1 forms, phospho-Akt^(Ser473)^ relative protein levels (phospho-Akt is normalized to Akt) and relative Krox20 mRNA and protein levels. Each experiment was repeated three times on different pools of three nerves per genotype. * P < 0.05 by one-way Anova with Bonferroni’s multiple comparison test. **(C)** Semithin cross-sections showing demyelinated axon (black arrow) in a 6 month old S63del sciatic nerve and quantification from at least three mice per genotype (lower histograms). * P < 0.05 by one-way Anova with Bonferroni’s multiple comparison test. **(D)** Semithin cross sections of 12 months old sciatic nerves: in both HANI and HANI/+//S63del mice myelin appeared properly compacted. Hypomyelination is still visibly rescued in small and medium caliber axons (arrowheads) in HANI/+//S63del mice as compared to S63del controls (size bar 25μm).

**Supplementary Figure 4.**
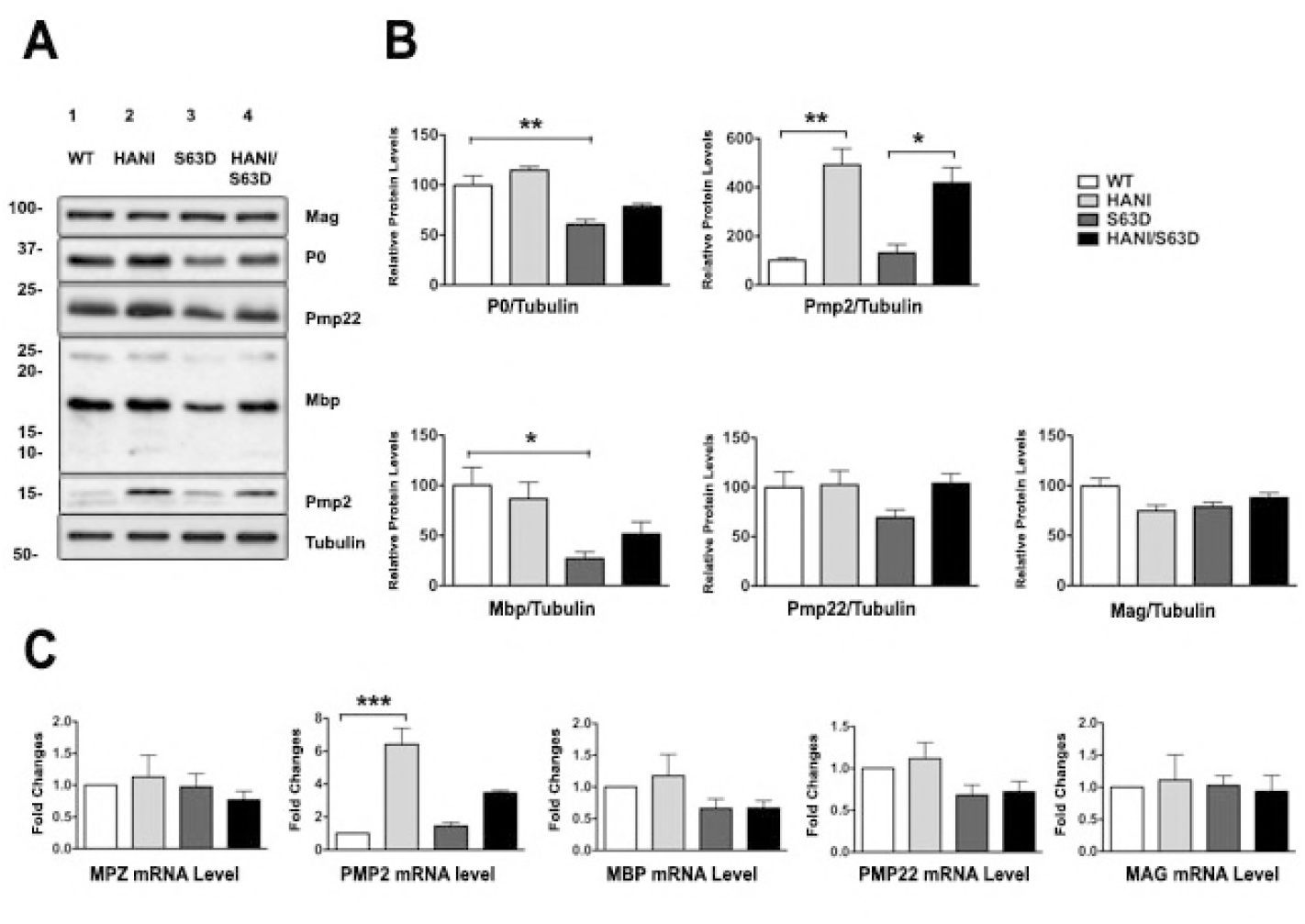
**(A)** Western blot analysis of 6 months old sciatic nerves lysates shows no differences in major myelin proteins expression between WT mice and those overexpressing Nrg1TIII, whereas the expression of P0 and Pmp22 is slightly, but not significantly increased in HANI/+//S63del as compared to S63del mice. Pmp2 protein expression is still significantly increased in HANI/+//S63del and HANI/+ mice as compared to S63del and WT nerves, respectively. **(B)** Quantification of P0, Pmp2, Mbp, Pmp22 and Mag relative protein levels. **(C)** Relative mRNA levels measured from sciatic nerves of P28 HANI/+//S63del mice and controls by qRT-PCR. No significant differences in expression for the major myelin genes analyzed (MPZ, MBP, PMP22, MAG) were detected among the genotypes, whereas there was a significant increase in PMP2 mRNA in HANI/+//S63del and HANI/+ mice as compared to S63del and WT mice, respectively. * P < 0.05, ** P < 0.01, *** P < 0.001 with One-way Anova with Bonferroni’s multiple comparison test. Each experiment was repeated three times on different pools of three nerves per genotype.

**Supplementary Figure 5.**
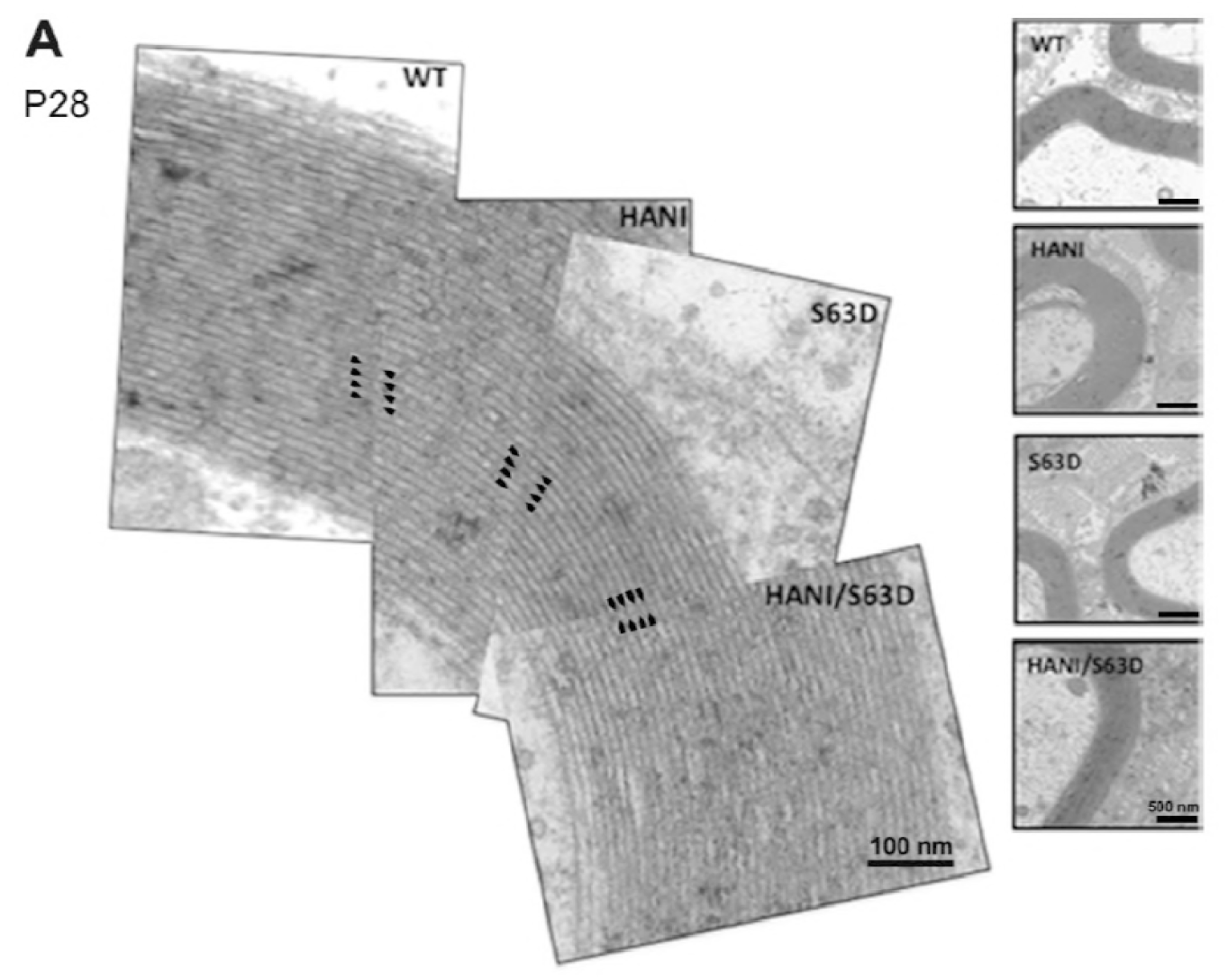
Comparison of myelin periodicity on electron microscopy images of cross sections of P28 sciatic nerves. There are no significant differences among the genotypes (see black arrows at equal distance), suggesting that myelin is normally compacted (size bar 100 nm). Images from at least three mice per genotype were analyzed.

**Supplementary Figure 6.**
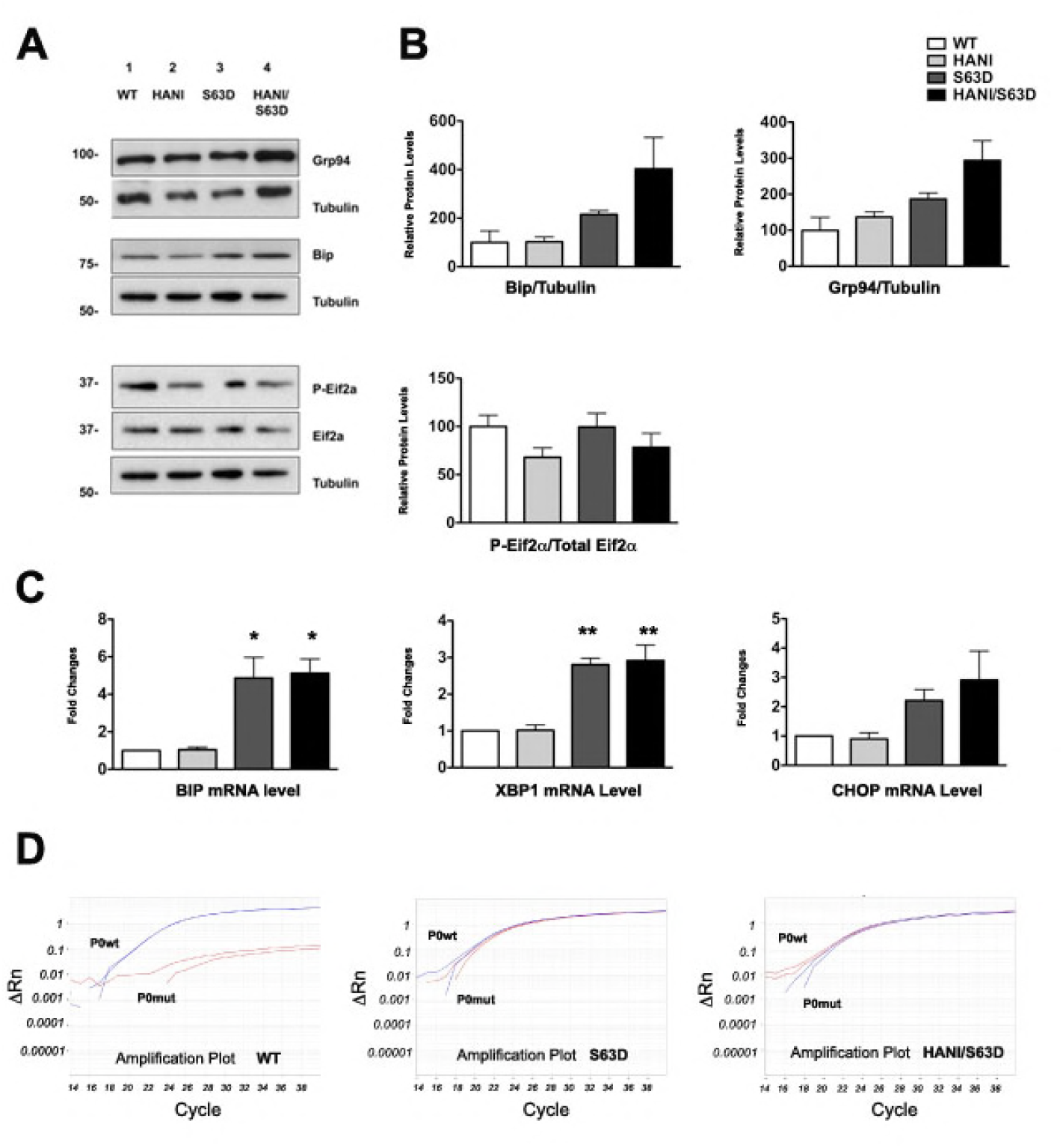
**(A)** Western blot analysis on 6 months old sciatic nerves lysates shows a slight, but not significant increase of ER stress (Bip and Grp94) and UPR (P-eIF2-alpha) markers expression levels in S63del mice as compared to WT and in S63del as compared to HANI/+//S63del mice. **(B)** Quantification of Bip, Grp94 and phospho-eIF2-alpha relative protein levels (phospho-eIF2-alpha is normalized to total eIF2-alpha). **(C)** mRNA levels measured by qRT-PCR for the UPR markers BiP, CHOP, and spliced XBP-1 in sciatic nerves at P28. In S63del nerves these factors are increased as compared to WT. No differences in the levels of expression were detected between S63del and HANI/+//S63del mice. **(D)** Allelic discrimination qRT-PCR assay shows no differences in the expression levels of wild-type and mutant P0S63del in presence of Nrg1TIII overexpression. * P < 0.05, ** P < 0.01 by One-way Anova with Bonferroni’s multiple comparison test. Each experiment was repeated three times on different pools of three nerves per genotype

**Supplementary Figure 7.**
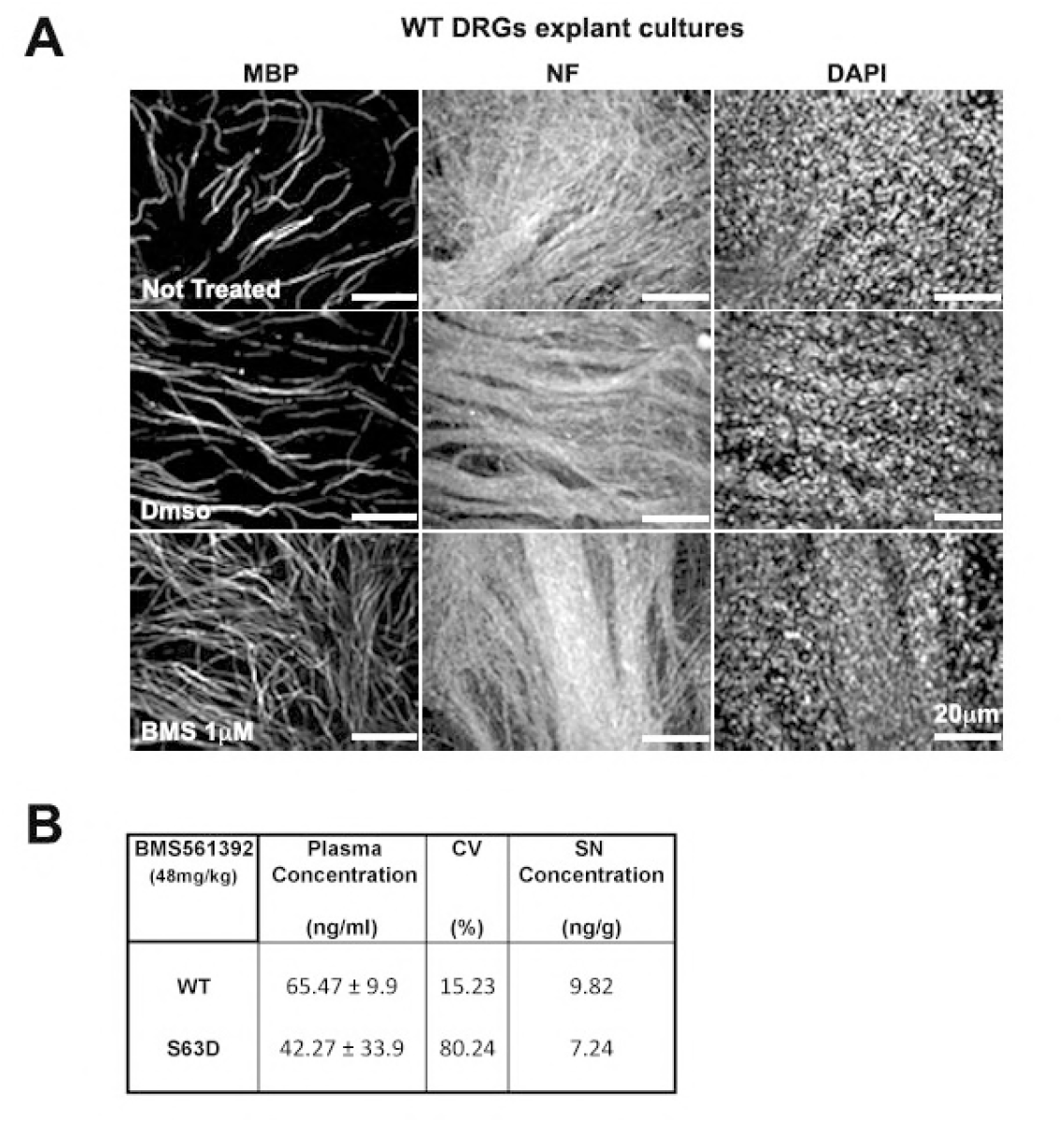
**(A)** WT myelinating DRGs treated with 1uM BMS for 2 weeks. As controls, untreated DRGs and DMSO-treated DRGs were analyzed. MBP staining shows an increase of MBP-positive segments in treated samples (size bar 20μm). For the quantification of WT internodes see Fig. 7B. **(B)** Bioavailability analysis of BMS levels in plasma and sciatic nerve after 10 days of *in-vivo* treatment performed twice a day via IP injection (at 48mg/kg); n=3 mice per genotype at 2 months of age. Data represent the mean ± SEM.

## Abbreviations

PNS: – Peripheral Nervous System
CMT1B: – Charcot Marie Tooth disease type 1B
MPZ (P0): – Myelin Protein Zero
P0S63del: – Serine 63 deletion at the N-terminal of P0
Nrg1TIII: – Neuregulin 1 Type III
HANI: – Nrg1TIII fuse to a N-terminal HA-tag
UPR: – Unfolded Protein Response
NCV: – Nerve Conduction Velocity
FWL: – F-Wave Length

